# Receptive field sizes and neuronal encoding bandwidth are constrained by axonal conduction delays

**DOI:** 10.1101/2023.01.15.524163

**Authors:** Tim Hladnik, Jan Grewe

**Author notes:** **Corresponding author:** Jan Grewe. **Author Contributions:** TH and JG designed the experiments, TH recorded the data, TH and JG analyzed the data, JG drafted the paper, TH and JG discussed and revised the paper.

## Abstract

Studies on population coding implicitly assume that spikes from the presynaptic population arrive simultaneously at the integrating neuron. In natural neuronal populations, this is usually not the case — neuronal signalling takes time and neuronal populations cover a certain space. The spread of spike arrival times depends on population size, cell density and axonal conduction velocity. We here analyze the consequences of population size and axonal conduction delays on the stimulus encoding performance in the electrosensory system of the electric fish *Apteronotus leptorhynchus*. We experimentally locate p-type electroreceptor afferents along the rostro-caudal body axis and relate locations to neurophysiological response properties. In an information-theoretical approach we analyze the coding performance in homogeneous and heterogeneous populations. As expected, the amount of information increases with population size and, on average, heterogeneous populations encode better than the average same-size homogeneous population if conduction delays are compensated for. The spread of neuronal conduction delays within a receptive field strongly degrades encoding of high-frequency stimulus components. Receptive field sizes typically found in the electrosensory lateral line lobe of *A. leptorhynchus* appear to be a good compromise between the spread of conduction delays and encoding performance. The limitations imposed by finite axonal conduction velocity are relevant for any converging network as is shown by model populations of LIF neurons. The bandwidth of natural stimuli and the maximum meaningful population sizes are constrained by conduction delays and may thus impact the optimal design of nervous systems.

**Author summary:** Reading out the activity of a population of neurons can yield a more complete picture of the stimulus space. Generally, increasing population sizes increases the amount of information. This applies to homogeneous populations of similarly tuned neurons as well as heterogeneous populations. So far studies on population coding assumed the information to reach the integrating neuron simultanously. But neurons are neither disembodied nor does neuronal signalling happen instantaneously; populations cover a certain space and axonal conduction takes time. The interplay of encoding bandwidth, receptive field respectively population size and axonal conduction velocity has not been addressed before. Our results from the electrosensory system and of model simulations show that stimulus dynamics, receptive field sizes, and conduction velocity have to be factored in when we want to understand the design of nervous systems.

## Introduction

Convergence is a basic network motif in nervous systems. Postsynaptic neurons integrate inputs from multiple presynaptic cells. Combining presynatic inputs can be beneficial by averaging out independent noise and increasing the information carried by the integrated response. This is, for example, observed in the retinae of vertebrates and invertebrates where neighboring photoreceptors are coupled to improve vision [1, 2]. If the input population is heterogeneous in the sense that different neurons encode different aspects of the stimulus space, the integrated response can provide a more complete representation of the stimulus space. Such a coding scheme is observed in color vision or olfaction in which the stimulus space is encoded in separate channels [3]. Heterogeneity can also refer to variations in the encoding properties within a given cell type [4, 5]. Modelling approaches have demonstrated that even in such cases the population response can carry more information about the stimulus than homogeneous populations of the same size [6–8].

Studies considering population coding usually combine presynaptic signals as arriving simultaneously at the integrating neuron. This is not necessarily the case since neuronal signalling is not instantaneous. Rather, information contained in the responses from one end of the receptive field will lead, while the action potentials originating from the other end will lag behind (Fig 1 A). The spread of conduction delays will depend on the receptive field size and the conduction velocity. Conduction velocity, in turn, depends on axon diameter or myolination and varies across modalities, tissues and species [9, 10]. Fast conduction is an investment that has to pay off and, hence, is mostly considered in the sense of minimizing delays and of mediating behaviorally important tasks such as escape responses [11, 12], when necessary for maximizing encoding bandwidth [10], or where the exact timing of a signal is essential for processing (e.g. in bird sound localization, [13]). The consequences of the spread of conduction delays within the population, has, to our knowledge, not been addressed so far.

**Fig 1.**
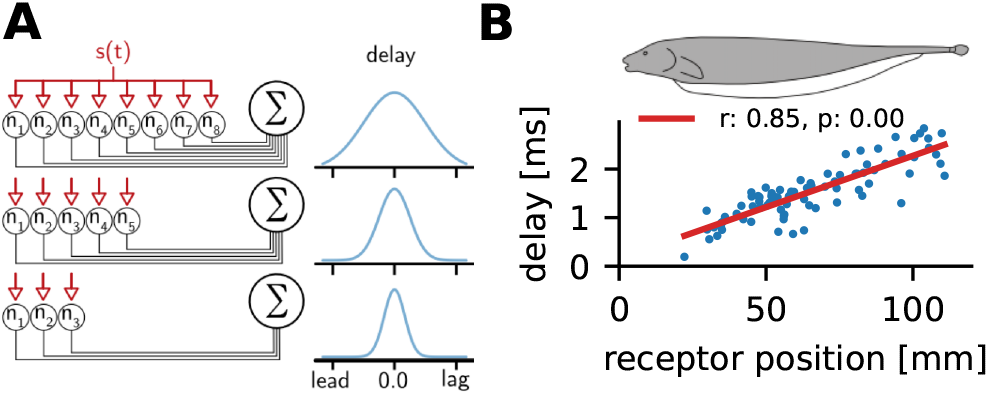
Spread of conduction delays depends on population size. A: Sketch of the integration of presynaptic inputs by a postsynaptic neuron. In this scenario presynaptic neurons are driven by the same stimulus (s(t)) and project to an integrating neuron. According to the spatial extent of the population, which is proportional to the size of the presynaptic population, the information from one end of the population will reach the integrating neuron earlier than from the other end. With respect to the center of the population, the first will lead while the latter will lag behind. B: The skin of the weakly electric fish *A. leptorhynchus* is a large sensory surface equipped with electroreceptors which project to integrating neurons in the hindbrain. The observed delay with respect to the stimulus depends on the location of the receptor on the body and is defined by the conduction velocity of the axons.

Here we investigate the interplay of population coding, heterogeneity, conduction delays, and encoding bandwidth in the active electrosensory system of the weakly electric fish *Apteronotus leptorhynchus*. Electric fish experience their electrical surroundings with two types of electrosensory subsystems, the active (tuberous) and passive (ampullary) system [14–17]. The active system and therein the population of p-type electroreceptor afferents have been extensively studied [18]. P-units are encoders of the fish’s electric field amplitude [19–22]. And their neurophysiological response properties are very heterogeneous with respect to their spontaneous as well as their stimulus driven activity [21–24]. About 8000 afferents per side of the fish [25] render the body a large sensory surface of heterogeneous and independent neurons [26, 27]. P-units project to the electrosensory lateral line lobe (ELL) in the hindbrain where they terminate in three adjacent segments of the ELL. The target segments differ in their spectral tuning as well as receptive field sizes [17, 28–30]. The delay with which spikes from electroreceptors arrive at the brain depends on the receptor location on the fish’s body (Fig 1 B). Hence, target neurons in the ELL will experience varying spreads of delays depending on their receptive field size.

In this study we first relate P-unit response properties and receptive field location along the rostro-caudal body axis. Then, in an information theoretical approach [31], we test whether the observed heterogeneity is indeed beneficial for the encoding of amplitude modulations and analyze the impact of population size and conduction delays on population coding. Finally, we compare and validate the above results to simulations using leaky integrate-and-fire (LIF) neuron models [32].

## Materials and methods

84 P-units were recorded in 13 subjects of *Apteronotus leptorhynchus* of either sex. Fish were of sizes in the range 10.5 to 22.3 cm and body weights in the range 4.1 to 14.6 g. P-type electroreceptor afferents were recorded in the posterior branch of the lateral line nerve. P-units were identified by their typical baseline firing properties and their response to amplitude modulations of the fish’s own field.

### Surgery

In a surgical intervention the lateral line nerve was exposed by opening a small patch of the skin just above the operculum where the nerve comes close to the surface. All experimental procedures complied with German and European regulations and were approved by the local district animal care committee (file no. ZP 1/16 Regierungspräsidium Tübingen).

Fish were initially anaesthetized with 125 mg/l MS-222 (PharmaQ, Fordingbridge, UK) dissolved in water taken from the housing tank and buffered to pH 7.0 using sodium bicarbonate. When deep anaesthesia was reached (gill movements ceased) fish were respirated with a constant flow of water through a mouth tube, containing 100 mg/l MS-222 (pH 7.0) to maintain anaesthesia during surgery. The lateral line nerve was exposed dorsal to the operculum. Fish were fixed in the setup with a plastic rod glued to the exposed skull bone. The wounds were locally anaesthetized with Lidocainehydrochloride 2 % (bela-pharm GmbH, Vechta, Germany) before exposing the nerve. Local anaesthesia was renewed every two hours by cutaneous application of Lidocaine to the skin surrounding the surgical wounds.

After surgery, fish were immobilized with 0.05 ml 5 mg/ml tubocurarine (Sigma - Aldrich, Steinheim, Germany) injected into the trunk muscles and were then transferred into the recording tank of the setup filled with water from the fish’s housing tank. Respiration was continued without anaesthetic. The animals were submerged into the water so that the exposed nerve was just above the water surface (see S1 Fig. Experimental setup). Electroreceptors located on the parts above water surface did not respond to electrical stimulation and were excluded from analysis. Water temperature was kept at 26 °C.

### Recording

Action potentials of electroreceptor afferents were recorded intracellularly with sharp borosilicate microelectrodes, pulled to a resistance between 50 and 70 MΩ when filled with a 1M KCl solution (Sutter P97 Brown Flaming type puller, Sutter Instruments, CA, USA). Electrodes were positioned by microdrives (Luigs-Neumann, Ratingen, Germany). The potential between the micropipette and the reference electrode (chlorided silver wire placed within the surgical wound close to the nerve) was 10x amplified and lowpass filtered at 10 kHz (SEC-05X, npi electronic GmbH, Tamm, Germany). Signals were digitized by a data acquisition board (PCI-6229, National Instruments, Austin TX, USA) at a sampling rate of 40 kHz (Signal No 1 in S1 Fig. Experimental setup). Spikes were detected and identified online based on the peak-detection algorithm [33].

The EOD of the fish was measured between the head and tail via two carbon rod electrodes (11 cm long, 8 mm diameter, signal No 2 in S1 Fig. Experimental setup, *Global EOD*). These electrodes were placed isopotential to the stimulation to ensure the recording of the unperturbed fish field. The transdermal potential was estimated by a measurement close to the skin by a pair of silver wire electrodes, spaced 1 cm apart, which were placed orthogonal to the side of the fish just behind the operculum (signal No 3, *Reference EOD*). The measurement of the reference EOD includes the fish’s field as well as the stimulus and is thus a proxy of the signal driving the electroreceptors. A third pair of silver wire electrodes were mounted on the arm of a XYZ-robot (mph automation, Reutlingen, Germany, signal No 4 in S1 Fig. Experimental setup, *Local EOD*) which could be moved alongside the fish.

EOD recordings were amplified by a factor between 100 and 1000 depending on the recorded fish’s EOD amplitude and bandpass filtered, cutoff frequencies of 3 Hz and 1.5 kHz for high and low-pass filter, respectively (DPA-2FXM, npi-electronics, Tamm, Germany).

Spike and EOD detection, stimulus generation and attenuation, as well as pre-analysis of the data were performed online during the experiment within the RELACS software version 0.9.7 using the efish plugin-set (www.relacs.net). Data was stored for further offline analysis in the open NIX data format (Resource ID: RRID:SCR_016196, https://github.com/g-node/nix, [34]). Datasets and all analysis code are available under the Creative Commons Attribution Non-Commercial Share-Alike 4.0 International License https://gin.g-node.org/jgrewe/hladnik_grewe_heterogeneity **Note: The repository will be frozen and receive an DOI upon acceptance of the paper**.

### Stimulation

The fish were stimulated either with artificial electric fields mimicking the presence of other weakly electric fish or with random amplitude modulations. Stimuli could either be global or local. In both cases stimuli were attenuated (ATN-01M, npi-electronics, Tamm, Germany), and isolated from ground (ISO-02V, npi-electronics, Tamm, Germany). Global stimuli were delivered via two carbon rod electrodes (30 cm length, 8 mm diameter) placed on either side of the fish parallel to its longitudinal axis. Stimuli were calibrated to evoke amplitude modulations of defined contrasts relative to the fish’s own unperturbed field measured close to the fish. Local stimuli were given using a pair of silver wire electrodes mounted on the xyz-robot (signal 6, S1 Fig. Experimental setup). Local stimuli could not be calibrated to the respective local conditions but stimulus intensities were increased until a noticeable response modulation was observed in the recorded P-units. The estimated receptive field positions are not affected by the stimulus intensity since relative changes are important for localization.

Random amplitude modulations (RAM) were created by multiplying the unperturbed global EOD with the desired noise waveform (frozen band-limited Gaussian noise, 0 – 300 Hz) and passing this signal into the experimental tank where it adds to fish’s EOD to result in the desired effect. The noise stimulus is calibrated to induce amplitude modulations with a standard deviation of 10% of the fish’s unperturbed field amplitude. RAM stimuli were always global.

### Data Analysis

Offline analyses were performed with custom routines in python 3.8 using the following open source packages: scipy [35], numpy [36], matplotlib [37], nixio [34], and pandas [38].

### Estimation of receptor position

The position of a recorded electroreceptor afferent was estimated by applying a local stimulus that was moved along the rostro-caudal axis by means of a local dipole electrode mounted on an xyz-robot (S1 Fig. Experimental setup). The receptor position was assumed to be at the position that led to the highest power at the expected frequency. The power spectrum was calculated from the time resolved firing rate of the neuronal response (see below) at any given stimulus position (S2 Fig. Estimating receptor position). A segment length of 1 s was used with 50% overlap. A Hanning window was applied to each data segment and means were subtracted. Cells were recorded in the posterior branch of the lateral line nerve. We could therefore only record neurons with receptive fields on the trunk of the fish.

### Baseline analysis

Each recording included several seconds of baseline activity in the absence of an external stimulus with only the fish’s own EOD being present (Fig 2A). Nevertheless, P-units show baseline firing that is highly irregular, sometimes bursty, but phase-locked to the EOD (fig. 2 A - C). To characterize the spontaneous activity, the baseline firing rate, the vector strength, the coefficient of variation of the interspike-interval (*CV_ISI_*) distribution and the burstiness were quantified. Spike time locking to the EOD carrier was evaluated using the vector strength [39, 40].

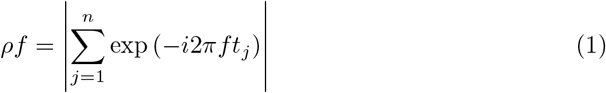

with *n* the number of spikes, *t_j_* the relative spike time of the j-th spike within the respective EOD cycle, and *i* the imaginary unit. The relative spike time is the delay between the spike time and the preceding rising threshold crossing at half-maximum of the EOD waveform. A vector strength of 0 indicates that spike phases are symmetrically distributed within the EOD period while a vector strength of 1 indicates perfect locking with all spikes occurring in the same phase of the EOD period.

**Fig 2.**
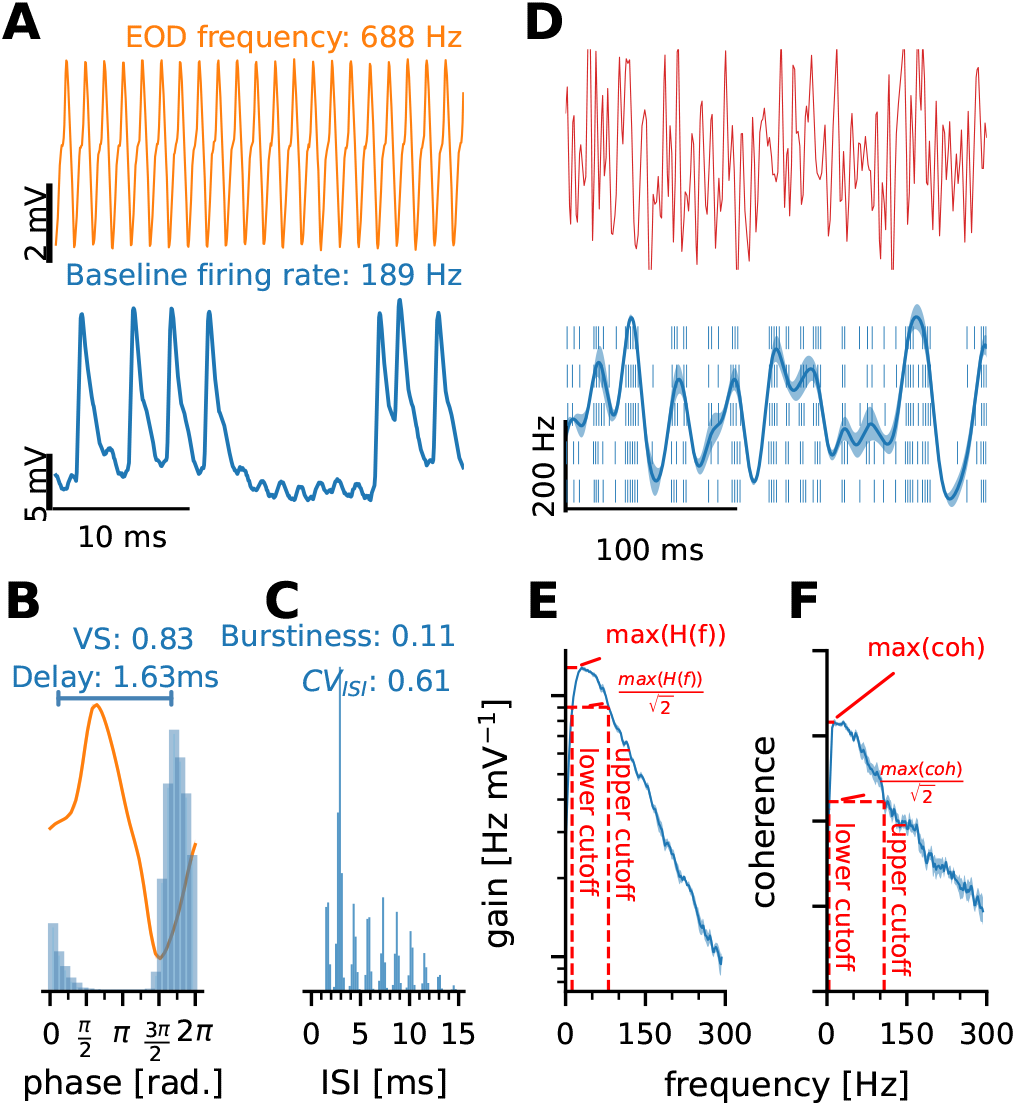
Analyzes on baseline and stimulus driven responses. **A - C:** Baseline response of an example P-unit. **A:** Short segments of the fish’s self-generated EOD (top) and the simultaneously recorded neuronal activity of a P-unit (bottom) in the absence of an external stimulus. This example neuron had a spontaneous activity of 189Hz. **B:** Spike times are precisely timed within the EOD period. The average EOD waveform is depicted in orange and the blue histogram shows the phases in which action potentials occur. From this we calculated the vector strength (VS, eq.1) which is 0.83 for this neuron. From the circular mean of the locking histogram we calculated the delay between EOD onset (time of the rising zero-crossing) and spiking activity. **C:** P-units skip EOD-cycles and sometimes fire bursts of action potentials. The interspike interval histogram shows the typical wide and multimodal distribution. Response regularity is characterized by the coefficient of variation (Eq 2, 0.61 in this example). The burstiness indicates the proportion of spikes occurring in intervals less than 1.5 times the EOD period [41] (0.11 in this example). **D: – F:** Driven responses of the same P-unit. **D:** The responses to frozen sequences of band-limited white noise of EOD amplitude modulations (0 – 300 Hz, top trace) are used to characterize the stimulus encoding properties. The stimulus profile is encoded in the spiking activity of the neuron (raster plot for 5 consecutive trials and firing rate in blue, bottom). The firing rate modulates around the baseline firing rate and the cell’s sensitivity is characterized by the strength of the modulation (Eq 4). The shaded area around the mean depicts the response variability, i.e. the across trial standard deviation of the firing rate. **E:** From the transfer function (Eq 6) we estimate the maximum gain as well as the upper and lower cutoff frequencies (red dashed vertical lines). **F:** Similar measures were extracted from the stimulus response coherence (Eq 7). The integral of the coherence spectrum is used a lower bound estimation of the mutual information between stimulus and response (Eq 8). The spectra in E and F have been smoothed with an 11 point running average. Abbreviations: VS, vector strength; CV, coefficient of variation; ISI, interspike interval; coh, coherence

The burstiness of the recorded cells was assessed using the burst fraction as proposed previously [41], i.e. the fraction of interspike intervals shorter than 1.5 times the average EOD period.

The coefficient of variation (*CV_ISI_*) is used to describe the response regularity [21, 22, 42]. A *CV_ISI_* of zero indicates perfect regularity of the response, a value of unity is characteristic of Poisson firing. The *CV_ISI_* is defined as:

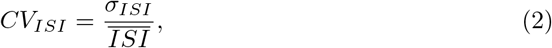

with *σ_ISI_* the standard deviation of the interspike-interval distribution and *ISI* the average interspike interval.

### Analysis of driven response

Characterization of the neuronal encoding of dynamic stimuli is based on the neuronal responses to band-limited white noise amplitude modulation stimuli (see stimulation, above; Fig. 2 D).

### Firing rate

The single trial firing rate, *y_k_*(*t*), was estimated by convolving spike responses with a Gaussian kernel

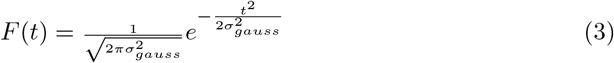

with *σ_gauss_* the standard deviation of the kernel which was 1 ms if not otherwise stated. The average firing rate (*y*(*t*)) is then calculated by averaging across trials.

### Estimating response modulation and variability

When driven by a stimulus the firing rate is modulated around the temporal average of the firing rate, 〈*y*(*t*)〉_*t*_, which is close to the baseline firing rate of the cell. The depth of the firing rate modulation is quantified by the standard deviation of the firing rate over time

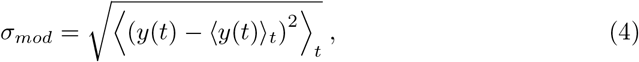

where 〈·〉_*t*_ denotes averaging over time. The response modulation is used here as a proxy of the cell’s susceptibility for the stimulus.

The response variability was quantified by the standard deviation of the single-trial firing rates *y_k_*(*t*) across trials averaged over time:

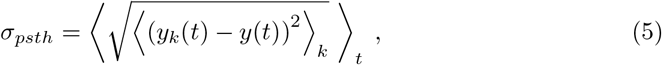

with *y*(*t*) the firing rate. Average firing rate and response variability are illustrated as solid line and shaded area in fig. 2 D.

### Spectral analyses

In addition to time domain analyzes we calculated the transfer function (*H*(*f*), Fig 2 E) according to:

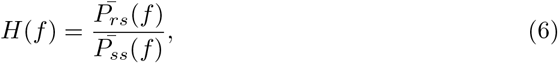

where 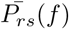 is the average cross spectral density of response and stimulus and 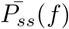 the average power spectral density of the stimulus. Averaging was done across trials and segments of approximately 1s duration (2^15^ samples) with 50% overlap. Segments were de-trended by subtracting the mean and a Hanning window was applied to each segment. From the transfer function we extracted the maximal gain as well as the lower and upper cutoff frequencies.

To estimate the (linear) information about the stimulus that is carried by the individual P-unit responses or by populations of P-units, the stimulus-response coherence 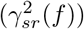 was calculated

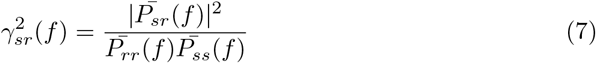

where 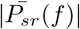 is the absolute of the cross spectrum and 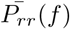 and 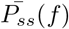 are the power spectra of the response and the stimulus, respectively. Averages are done across trials and segments. Spectra were estimated as described for the transfer function above. From the coherence spectrum we calculated the coherence rate, as a lower bound estimate of the mutual information between stimulus and response [43]

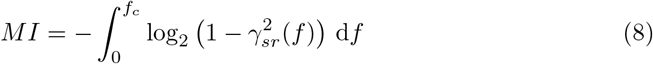

in which *f_c_* is the cutoff frequency of the stimulus (300 Hz).

### Artificial populations of P-units

The information about the stimulus that is carried by neuronal responses was further estimated for artificial homogeneous and heterogeneous populations. Since we did single unit recordings these populations were constructed from neurons recorded in different animals in different recording sessions. In addition to the neurons recorded within this project we added 70 P-units recorded in a different project [22]. In total, the population analyses are thus based on 130 neurons recorded in 36 fish. All cells were stimulated with the same frozen Gaussian white noise AM stimulus.

From each neuron a set of up to 10 homogeneous populations was created for population sizes ranging from 1 to the number of trials recorded in that particular neuron. Population responses were assembled from unique random combinations of trials. The population response is the average firing rate estimated with a Gaussian kernel of 1.0 ms standard deviation. From each population response the response modulation (Eq 4) and the average firing rate were extracted. Additionally, the stimulus response coherence was estimated (Eq 7) and the mutual information (Eq 8) as well as the lower and upper cutoff frequencies of the coherence spectrum were estimated.

Heterogeneous populations were created by randomly drawing trials from all recorded neurons. Per population each cell could contribute with at maximum one trial. For each population size (2 to 30 neurons) 50 different populations were created. Each selected trial was then temporally aligned with the stimulus by subtracting the average delay (estimated from the cross-correlation of stimulus and response) from the recorded spike times. The population response was estimated as the average PSTH across trials. When artificial delays were added, all spikes of an individual trial were shifted by the same delay that was drawn from a Gaussian distribution with zero mean and the respective standard deviation (*σ_delay_*) after the response was temporally aligned to the stimulus as described above.

### Leaky integrate and fire model

Leaky Integrate and Fire (LIF) [32] model neurons were created by numerically solving

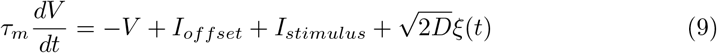

in which *τ_m_* is the membrane time constant, *V* the membrane voltage, *I_offset_* an offset current, *I_stimulus_* the stimulus current and an additional noise current drawn from a Gaussian distribution *ξ*. Δ*t* is the temporal stepsize of the simulation. Whenever the membrane voltage exceeded the threshold *θ*, the spike time was noted, and the membrane voltage was reset to *V_reset_*.

The population was modelled as a 1-dimensional array of LIF model neurons with identical parameterization and independent noise. On the one-dimensional array spacing of neurons was assumed to be equal. All model neurons were driven with the same 2s frozen white noise sequences which had spectral power in the ranges 0 – 100 Hz, 100 – 200 Hz, or 200 – 300 Hz created from Gaussian white noise (strength of the stimulus is controlled via the standard deviation, *σ_stimulus_*). All model parameters are listed in table 1. The population response was estimated as the average firing rate estimated by kernel convolution as described above using a Gaussian kernel with a standard deviation of 1.25 ms.

**Table 1.** Model parameterization. Parameters in the membrane model part refer to eq 9

| Membrane model |  |  |  |  |  | Population model |  |
| --- | --- | --- | --- | --- | --- | --- | --- |
| $\tau_m$ | $I_{offset}$ | $\sqrt{2D}$ | $\theta$ | $V_{reset}$ | $\Delta t$ | $\sigma_{stimulus}$ | cell density |
| 0.25 ms | 2.5 | 5 | 1 | 0 | 0.01 ms | 0.0625 | 2000 m<sup>-1</sup> |

## Results

A total of 84 p-type electroreceptor afferents were recorded in 13 subjects of *Apteronotus leptorhynchus* to estimate the population heterogeneity and its potential dependency on the receptor’s position. In each cell we recorded the baseline activity and, in a subset, the responses to dynamically varying sequences of frozen band-limited white noise amplitude modulations. From baseline and driven responses we extracted several features to characterize the cells (Fig 2).

### Most baseline parameters are not correlated with receptive field position

The majority of the recorded P-units had receptive field positions between 30 and 50% of the total body length (Fig 3 A). In our sample no cells with receptive field positions beyond 80% of the body length could be recorded. Cells more frontal than about 20% of the body length could not be recorded at the recording location (see methods). In the absence of an external stimulus (but in the presence of the unperturbed own electric field) P-units show a spontaneous baseline activity (Figs 2 A-C) that is very heterogeneous across cells, even when recorded in the same animal [21, 22]. From the baseline activity we extracted five characteristics: the baseline firing rate, the coefficient of variation of the interspike intervals (*CV_ISI_*), the fraction of spikes that occurred in bursts of action potentials, and from the locking of the spike times to the EOD we extracted the vector strength and the preferred phase (see methods, Fig 2 A-C). These features were then correlated with the estimated receptor position (Fig 3 B – F). The overall range of observed firing rates varies between about 60 and 530 Hz (249±117 Hz) matching previously published observations [21, 22]. The *CV_ISI_* (0.52±0.19), the burst fraction (0.3±0.26), and the vector strength (0.86±0.05) fall well into the ranges reported before. The firing rate, *CV_ISI_* and vector strength do not statistically significantly depend on the receptor’s position (Fig 3 B, C, E). The burst fraction shows a statistically significant decline in our sample (r=-0.30, p=0.02). The vast spread of the data and the few points at the most frontal and most caudal positions, may bias this observation. The phase relation between the spike times and the fish’s own electric field, however, depends clearly on the receptor position (r=0.75, p < 0.01, Fig 3F).

**Fig 3.**
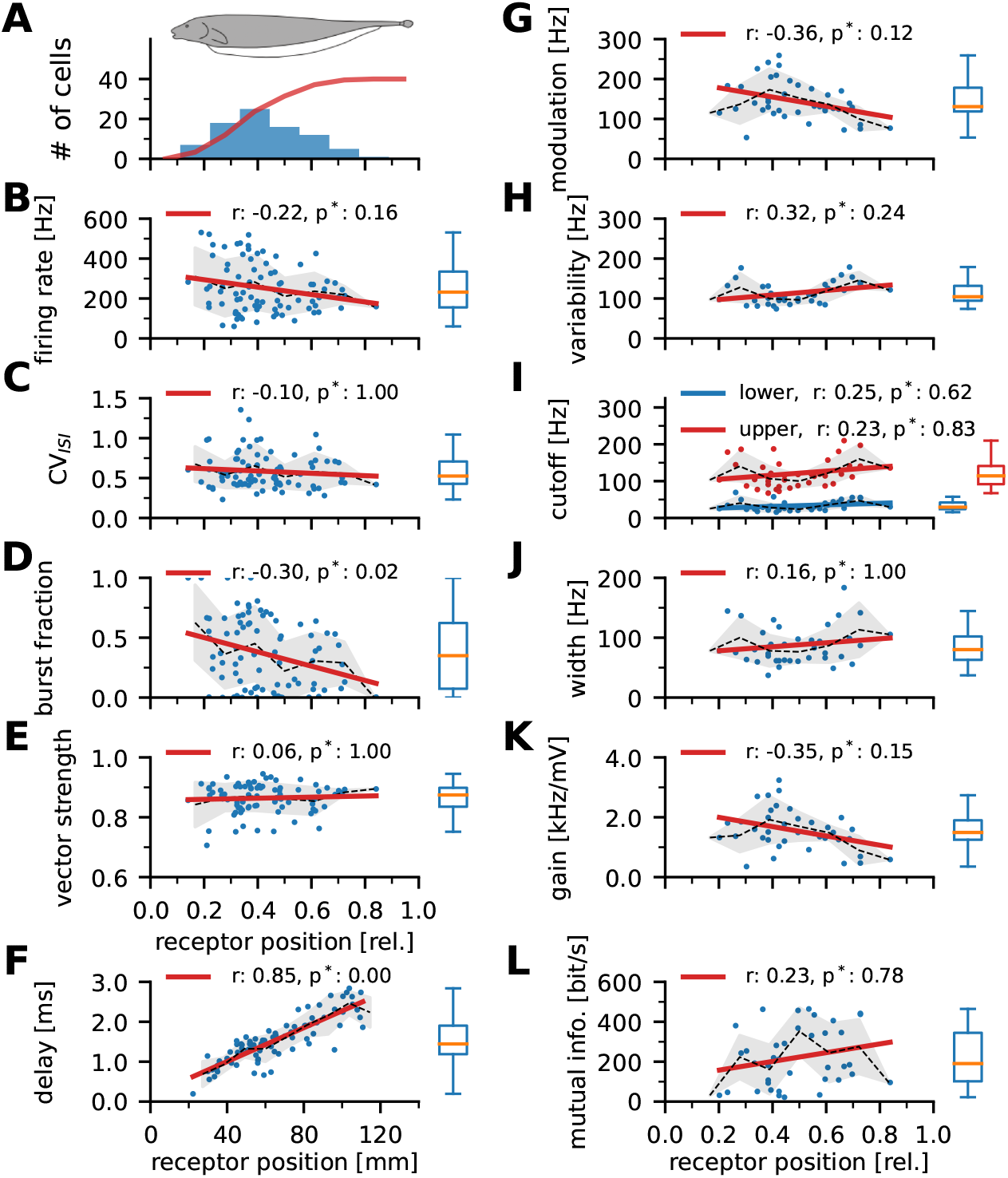
Correlation of response features and receptor position. r-values are the Pearson correlation coefficients, p-values are Bonferroni corrected. Boxplots show the spread of the data, whiskers depict 1.5 of the inter-quartile range. **B – F**: properties of the baseline activity, **G – L:** characteristics of the encoding of dynamic stimuli. **A:** In this study 84 p-type electroreceptor afferents were recorded. The blue histogram shows the distribution of receptor positions along the rostro-caudal body axis expressed relative to the total body length. The cumulative (red line) indicates that three fourths of the recorded cells were recorded in the frontal third of the body. Due to the recording location cells in the frontal 20% of the body could not be recorded. **B:** Baseline firing rate estimated with only the unperturbed fish’s field present. **C:** Coefficient of variation of the interspike interval distribution (*CV_ISI_*) during baseline activity. **D:** Fraction of action potentials observed as part of action potential bursts as previously defined [41]. **E:** Vector strength quantifying the phase-locking to the EOD. **F:** Delay between the preferred phase in the EOD cycle and the beginning of the EOD. **G:** Response modulation of the across-trial average firing rate (Eq 4) in response to frozen white noise amplitude modulations. Please note that these stimuli were only presented to a subset of 39 neurons which could be recorded sufficiently long to allow for position estimation and stimulation with white noise stimuli and were stimulated with the same stimulus intensity (10% contrast). **H:** response variability, i.e. the across-trial standard deviation of the firing rate (Eq 5). **I:** lower (blue) and upper (red) −3dB cutoff frequencies of the transfer function. **J:** Tuning width, i.e. the difference of the upper and lower cutoff frequency. **K:** peak gain of transfer function. **L:** Mutual information between responses and the stimulus estimated from the stimulus response coherence spectrum.

### Delay shift can be explained by axonal conduction velocity

The vector strengths depicted in Figs 2 B and Fig 3 E show the tight temporal coupling between P-unit spikes and the EOD period [40, 44]. The delay between EOD onset (time of the zero-crossing in the rising flank of the EOD, see methods for details) and the circular mean of the spike-times within the EOD cycle increases the more caudal the receptive field is. The data can be well-fitted with a linear regression and the inverse of the slope of the regression line is then the axonal conduction velocity. With this across-animal approach we estimated it to 48.3 m/s which matches to the within-animal estimation from the temporal differences in preferred phase from all possible pairs of P-units recorded in the same animal (see S3 Fig. Dependency of phase locking on receptor position D).

### Coding properties are not correlated with receptive field position

In a subset of 39 neurons recording durations allowed for analyzing the stimulus encoding performance using sequences of white noise amplitude modulations of the fish’s field (Fig 2 D-F, see methods). The cellular responses were characterized by the response modulation of the across-trial average firing rate (Eq 4, Fig 3 G, 145 ±49Hz), the across-trial response variability (Eq 5, Fig 3 H, 114±27Hz), and by using the transfer function to characterize the spectral coding range. From the transfer function we extracted the lower and upper cutoff frequency (33±13 Hz and 121±37 Hz, respectively), the width of the coding range (88±32Hz), and the maximum gain (1553 ±32 Hz/mV, Fig 3 I, J, and K, respectively). From the coherence spectrum we quantified the amount of information carried by the responses about the stimulus by estimating the lower-bound mutual information as the integral of the coherence spectrum (Eqs 7 and 8, [43]). In the population of recorded cells the mutual information varies strongly (Fig 3 L, median: 181 bit/s, interquartile range from 106 to 286 bit/s). All measures of the stimulus encoding performance vary considerably across cells, but none showed a statistically significant correlation with the receptor location.

### Pyramidal cells in the ELL might gain from population heterogeneity in their input layer

Postsynaptic neurons in the ELL receive information from up to about 1000 P-units [29]. Since most of the baseline as well as the stimulus encoding properties of the P-units do not depend on the location on the body (except for the preferred phase), each pyramidal neuron will integrate the activity of a similarly heterogeneous population of P-units.

To test whether stimulus encoding in pyramidal neurons profits from P-unit heterogeneity, we constructed artificial homogeneous or heterogeneous populations from the pool of recorded P-units (sell also S5 Fig. Stimulus encoding with homogeneous and heterogeneous populations). All cells were stimulated with the same white noise stimulus of amplitude modulations which has spectral power in the range 0 – 300 Hz. For the following analyzes we extended our dataset to 130 cells including those cells for which no receptor location was estimated (60 in total from this study) plus 70 cells that have been recorded in a previous study [22].

Homogeneous populations were constructed by combining responses recorded in the same neuron. Depending on the recording duration the number of recorded trials varies across cells and therefore the maximum population size varies accordingly. With an increasing population size the mutual information between the population response and the stimulus increases (blue lines in Fig 4 A). On average, the amount of information carried by the homogeneous populations roughly doubles from 1 to 30 cells (thick blue line, 261 bit/s with two cells to 536 bit/s with 30 cells).

**Fig 4.**
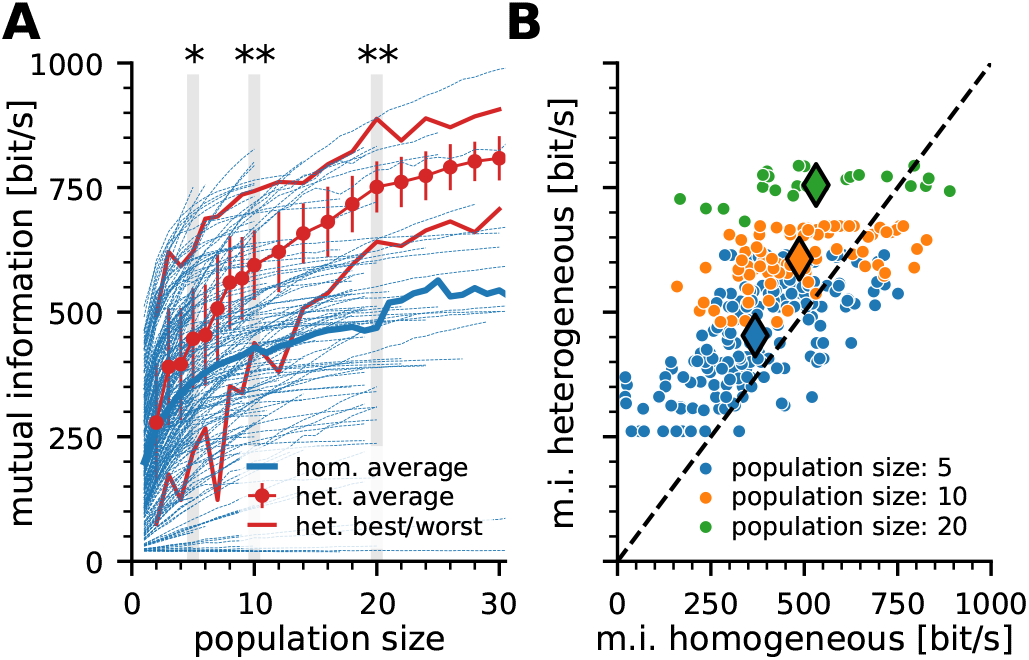
Information content of homogeneous and heterogeneous populations of P-units. **A:** Mutual information estimated from the stimulus response coherence (Eq 8) carried by populations of increasing size. Blue lines show information content of homogeneous populations created by combining randomly chosen trials of the same neuron. Since not all recordings have the same number of stimulus repetitions, the maximum possible population size varies between recorded neurons. Thick blue line is the average across all homogeneous populations. Red dots show the average information estimated from heterogeneous populations (see methods for details). Error bars indicate the standard deviation, thin red lines depict the best and worst performance observed in the sample of heterogeneous populations. Tests on statistically significant differences in mutual information in homogeneous and heterogeneous population were performed using a Mann Whitney U-Test with Bonferroni correction highlighted population sizes, one asterisk: *α* < 5% two stars indicate *α* < 1%. **B:** Comparison of the mutual information carried by homogeneous (x-axis) and heterogeneous (y-axis) populations that are similarly driven by the stimulus (response modulation of heterogeneous population response within a ±20% range of the respective homogeneous response) for three different population sizes. Large symbols depict the center of gravity of the respective distributions. Dashed line is the identity line. Abbreviations: m.i., mutual information; het., heterogeneous; hom., homogeneous

Heterogeneous populations were constructed from combinations of responses from different cells that may even be recorded in different animals (see methods for details). The population response was calculated after the individual responses were temporally aligned with the stimulus which eliminates variations in delays. As for homogeneous populations the information carried by heterogeneous populations increases with population size (295 bit/s with 2 cells to 704 bit/s with 30 cells). The average heterogeneous population carries more information about the stimulus than the average homogeneous population. The best heterogeneous population does not exceed the performance of the best homogeneous populations. But, conversely, the worst heterogeneous population performs generally better than many homogeneous populations (Fig 4 A, red dots, thin red lines for best and worst heterogeneous population). The information gain through heterogeneity is statistically significant for populations larger than four cells (tested for the highlighted population sizes in Fig 4 A). In general, the firing rate modulation is a good predictor of the mutual information (S4 Fig. Correlations of cellular response properties). For a fair comparison we compare the coding performance of homogeneous and heterogeneous populations that show similar (± 20%) response modulations of the population response (Fig 4 B). For populations larger than five cells, the vast majority of heterogeneous populations carries more information than the respective homogeneous ones. The center of gravity deviates increasingly from the identity line (large diamond markers) for larger populations.

### Neuronal conduction delay acts as an information filter in the population response

In the population analysis described above the responses were aligned to the stimulus and thus every spike contributing to the population had the same delay. This is not the case in real populations when axon lengths vary depending on their origin within the population (Fig 3 F).

The heterogeneous populations were now used to systematically study the effect of conduction delays on stimulus encoding performance. As above, responses were aligned to the stimulus and then an artificial delay (drawn from a Gaussian normal distribution with varying standard deviation, *σ_delay_*, assuming consistent conduction velocities and receptive fields as was previously described [29]) was added to the spikes. The same delay was applied to all spikes originating from the same neuron to mimic conduction delays and not to increase spike time jitter.

With increasing *σ_delay_* the amount of information drops. The larger the delay, the more severe is the effect. In the extreme, there is hardly any gain through population averaging (Fig 5 A, increasing *σ_delay_* from light to dark color). When separating the stimulus encoding performance for three different spectral bands (Fig 5 B) it becomes clear that the deteriorative effect is not equally strong for all stimulus components. Low-frequency components are more robust against delays in the population than high-frequency components. Independent of the added delay, P-unit populations encode higher frequencies worse than lower spectral bands (compare also to the transfer or coherence spectra shown in Fig 2 E, F). Encoding high-frequency changes in the stimulus requires higher precision in the spike times than encoding slow changes and is thus more sensitive to delays across the population (Fig 5 C for a population size of 20 neurons, normalized to the respective coding performance at zero delay).

**Fig 5.**
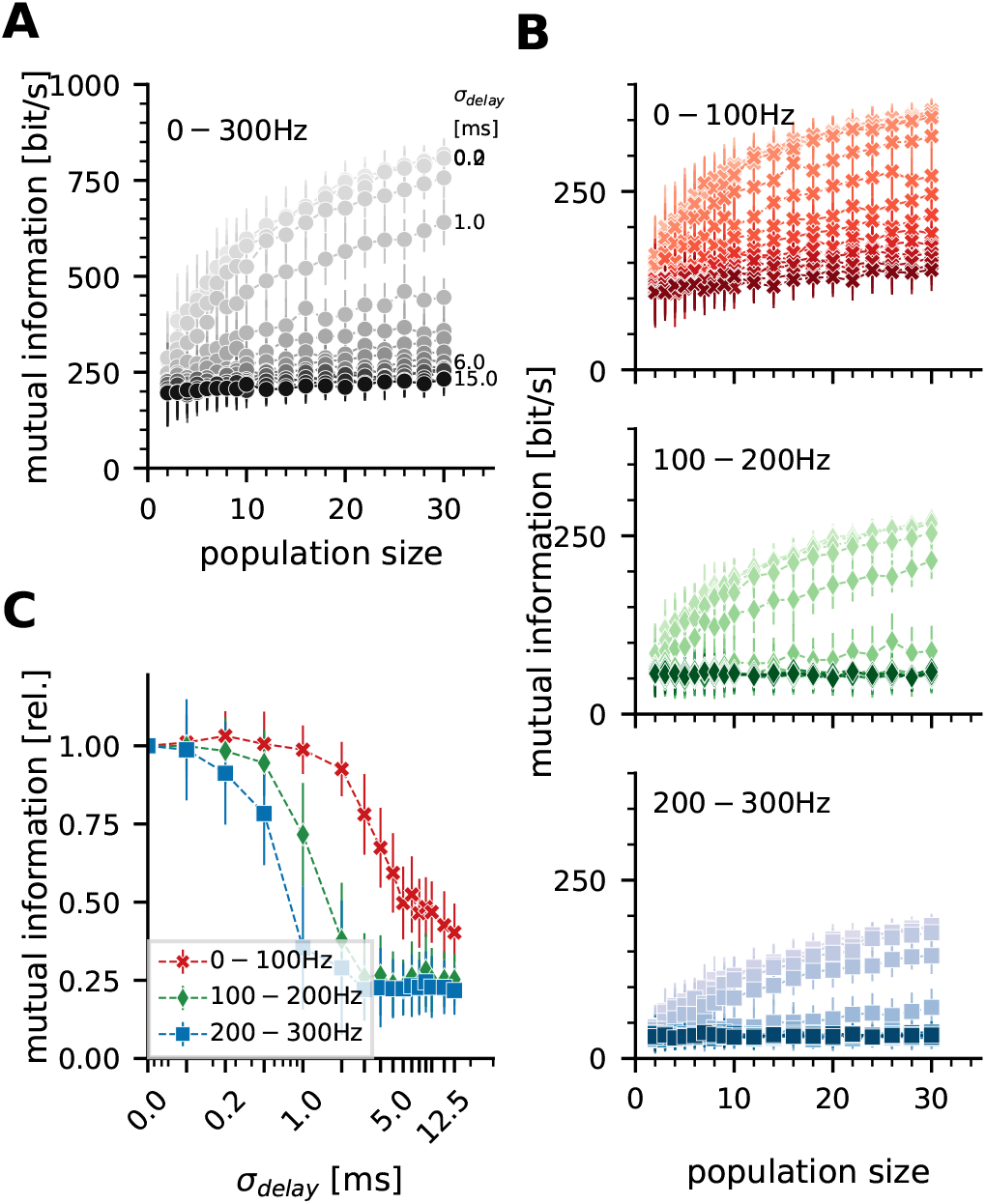
Conduction delays in a population act as an information filter. **A:** Average mutual information carried by heterogeneous populations of increasing population size that are subjected to varying simulated conduction delays. **B:** Mutual information as a function of population size for three spectral bands (low frequencies 0 – 100 Hz, red, top panel; intermediate 100 – 200 Hz, green, center panel; high 200 – 300 Hz. blue, bottom panel. Increasing opacity indicates increasing spread of conduction delays within the population as in A. **C:** Mutual information as a function of conduction delay compared for the three different spectral bands as in B. High frequency information is more strongly affected than low-frequency content.

### Conduction delays and stimulus dynamics constrain meaningful population sizes in populations of model neurons

The results shown above indicate that the amount of information conveyed by populations of P-units is affected not only by the population size and the noise in the system but also the spread of conduction delays within the population. In particular, high-frequency information suffers from a too large spread of conduction delays. Assuming a consistent conduction velocity and a homogeneous receptor density across the sensory surface, this is directly related to the spatial extent of the population. To extend these ideas beyond the electric fish, we performed simulations using a leaky integrate and fire (LIF) models with independent noise for each model neuron. The model neurons were not fitted to any realistic neuron (for details on the model parameterization see methods). The model cells were stimulated with band-limited white noise stimuli which had spectral power in three different ranges (0 – 100 Hz, 100 – 200 Hz, and 200 – 300 Hz) and project to an integrating neuron via axons with three different conduction velocities (7 m/s as observed in visual fibers in the monkey corpus callosum [45], 25 m/s as a representative for the squid giant axon [9], and 50 m/s as estimated here for the p-type electroreceptor afferents). An arbitrary cell density of 2000 *m*^−1^ was assumed. The mutual information between the population response and the stimulus was again estimated from the stimulus-response coherence.

Increasing the population size increases the mutual information if no conduction delay is assumed (blue solid lines in Fig 6 A, B, and C). When considering realistic conduction velocities, the arising spread of conduction delays within the population (distance between neuron *n*_1_ and *n*_8_ divided by conduction velocity Fig 1 A) affects the amount of information carried by the population response. In line with the previous experimental observations, the severity depends on the stimulus dynamics and the conduction velocity (broken, dash-dotted and dotted lines representing different stimulus dynamics in Fig 6 A, B, and C). The lower the conduction velocity, the stronger the limitation through delays, respectively population size, respectively spatial extent. For the lowest velocity used here and the highest frequency band (7m/s, 200 – 300 Hz, Fig 6 A, dotted line) increasing the population size reduces the mutual information down to the single-neuron performance. The ‘‘harmonic” structure reflects the autocorrelation of the stimulus. The faster the axonal conduction velocity, the more robust is the system and the larger the populations may be. Hence, faster conduction allows for larger population sizes and a higher possible gain from the integration over large populations.

**Fig 6.**
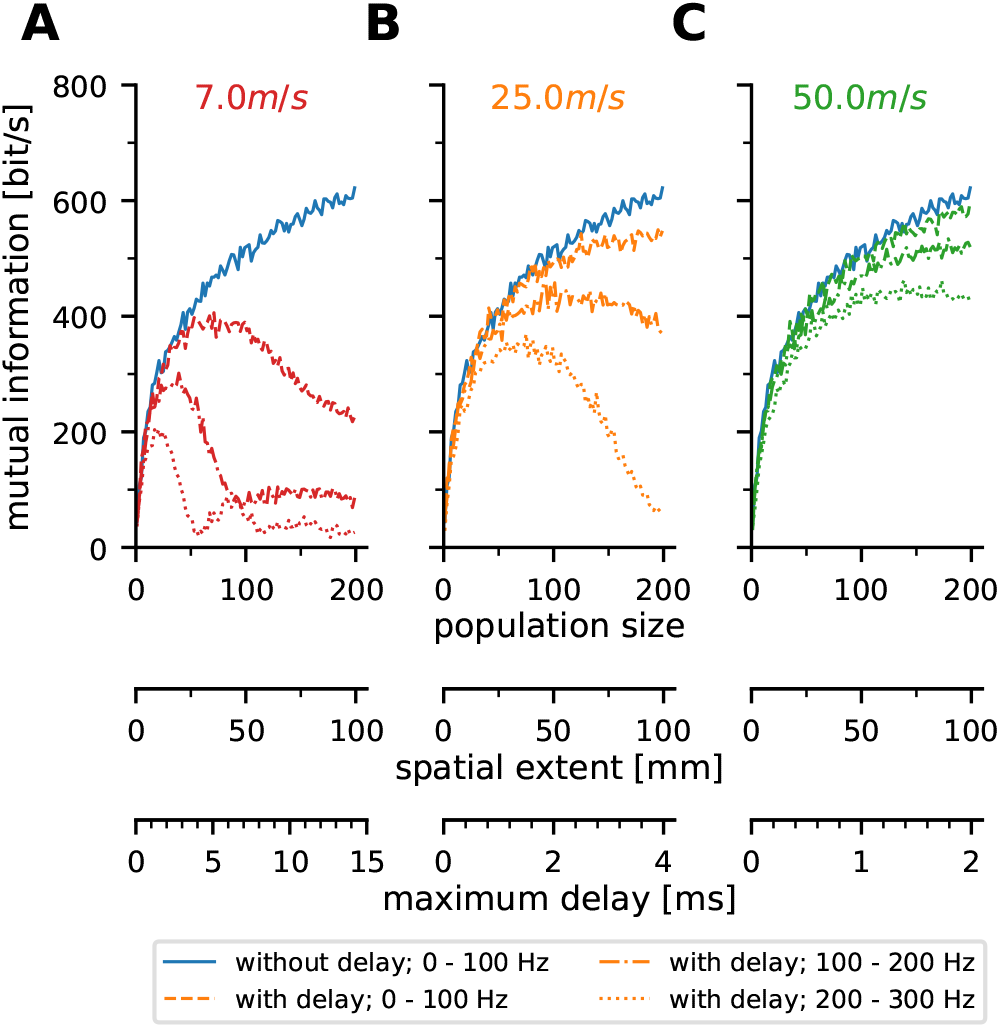
Conduction velocity and stimulus dynamics constrain meaningful population sizes of LIF model neurons. **A - C** Mutual information between stimulus and response estimated from the population responses of LIF-model neurons (see sketches in Fig 1 A). These toy-model assumes 1D populations with a density of 2000 neurons per m of the sensory surface. Model parameterization was completely arbitrary and identical for all model neurons (see methods). The models were driven with the same frozen noise sequences with spectral power in the ranges 0 – 100 Hz, 100 – 200Hz and 200 – 300Hz (dashed, dash-dotted, and dotted lines, respectively). Each model had the same amount of (independent) noise added to the driving stimulus. Three realistic conduction velocities were simulated and compared to the unrealistic instantaneous conduction (blue solid line in all figures, ‘‘without delay”). With decreasing conduction velocity the encoding performance drops at smaller populations. High-frequency encoding is most sensitive to spread in conduction delays. The ‘harmonic” structure seen in A is a consequence of the stimulus’ autocorrelation. **A:** Conduction velocity of 7 *ms*^−1^, as described for some visual fibers in the monkey corpus callosum [45] **B:** 25 ms^−1^ for example measured in squid giant axons [9], **C:** 50 ms^−1^ as estimated above for P-unit afferents.

## Discussion

Here we studied the heterogeneity among p-type electroreceptor afferents in *A. leptorhynchus*. The spatial homogeneity in P-unit heterogeneity implies that postsynaptic neurons in the hindbrain will always receive heterogeneous inputs from the sensory periphery. The system can profit from the heterogeneity since, on average, heterogeneous populations carry more information about the stimulus than the average homogeneous population of the same size. Generally, mutual information increases with population size but integrating over a spatially large patch of the sensory surface unavoidably leads to a spread of temporal delays within the population. This is an effective filter on the information content and particularly critical for high-frequency stimulus components. An effect that is not fish-specific but is also observed in populations of leaky integrate-and-fire model neurons.

### P-unit population is homogeneously heterogeneous

The presented data confirms previous observations of the large spread of baseline and stimulus encoding properties in P-units [21, 22, 24, 42]. Only minor parts of the observed variance can be explained by the receptor location. Even in the burst fraction, the only parameter found to statistically significantly depend on location, only 9% of the total variation can be attributed to the position (Fig 3D).

P-units are known to fire action potentials tightly phase-locked to the EOD which can be characterized by the vector strength [15, 39, 40, 46](Fig 2 B). The preferred phase of action potential firing within the EOD period (Fig 3F) shows a strong dependency on receptor location which can be fully explained by the distance between receptor location and recording site assuming a constant axonal conduction velocity. The conduction velocity was estimated between 48.2 m/s and 57.1 m/s which is faster than described for other Gymnotiform electric fish: For *Eigenmannia virescens* conduction velocities of 30 *m/s* for receptors on the trunk and 15m/s for afferents on the head were reported [47]. For *Hypopomus sp*. speeds of 25 m/s were reported but this study does not describe differences between frontal and caudal receptors [48]. Soma volumes of tuberous afferents in *E. virescens* were shown to be larger for more distant receptors and a correlation between soma volume and axon diameter and thus conduction velocity was suggested [47]. Our data does not show such dependence and also the original data seems to be well-fitted by a linear regression [47].

Conduction velocity depends on the axonal morphology, increasing it is an energetically expensive investment [10]. While both, *E. virescens* and *A. leptorhynchus* are wave-type electric fish, their EOD frequencies occupy very different ranges, 250 to 450Hz in *E. virescens* and 600 to 1000 Hz in *A. leptorhynchus* [49, 50]. This higher EOD frequency in *A. leptorhynchus* enables it to encode a wider signal bandwidth (as predicted from the sampling theorem), but in order to preserve the additional information higher conduction velocities are needed. From the *E. virescens* perspective investing in faster conduction may not pay off if the stimulus dynamics are limited to lower frequencies for other reasons.

### Origin of heterogeneity in P-units is unclear

Previous work shows that the tuning of newly generating electroreceptor organs is off for a few weeks [51], these organs are usually smaller with fewer primary electroreceptors per organ and there is variability in the number of electroreceptor organs innervated by a single afferent fiber [52]. Such variations may have consequences for the sensitivity and the response reliability of the P-unit in total. Geometrical effects due to the curved body surface [48, 53] and the fact that the electric field intensity is not uniform over the body surface [15] will have an impact on the cellular sensitivity as well but do not explain the heterogeneity in other parameters.

### Heterogeneity beneficial in more than one context

When two fish interact, the interference of their fields not only leads to regular amplitude modulation (AM) but, as the fish move relative to each other, the strength of the AM will change as a function of time [54]. Encoding such second-order AMs (also referred to as envelope) was shown to be facilitated by heterogeneity among P-units. Saturation non-linearities in neurons with particularly high or low firing rates enable the P-units to encode such signals [24]. Our population analyzes show that heterogeneity is beneficial for the encoding performance even when we only consider linear encoding (as the stimulus response coherence is a linear method). We would argue that even if some cells are saturated by the stimulus, others in the population will still be able to encode changes in stimulus amplitude. Thus, the interpretations are not mutually exclusive but hint at heterogeneity being beneficial in more than one context.

### Conduction delay as information filter

The spread of conduction delays within populations strongly affects the information carried by the population response. *σ_delay_* acts much like a low pass filter attenuating high-frequency content more effectively than low-frequency content. Representing high-frequency stimuli requires high spike time precision. Differences in spike arrival from the two ends of the population will inevitably destroy the required precision (Fig 1 A illustrates this schematically). Comparing the impact of temporal low-pass filtering as would happen at chemical synapses (termed *σ_kernel_*) and that of the delay (*σ_delay_*, Fig 7) one can observe that mutual information measures are higher above the identity line and the drop-off is steeper for *σ_delay_* than for *σ_kernal_*. The spread of conduction delays is more detrimental than the low-pass filtering at a chemical synapse.

**Fig 7.**
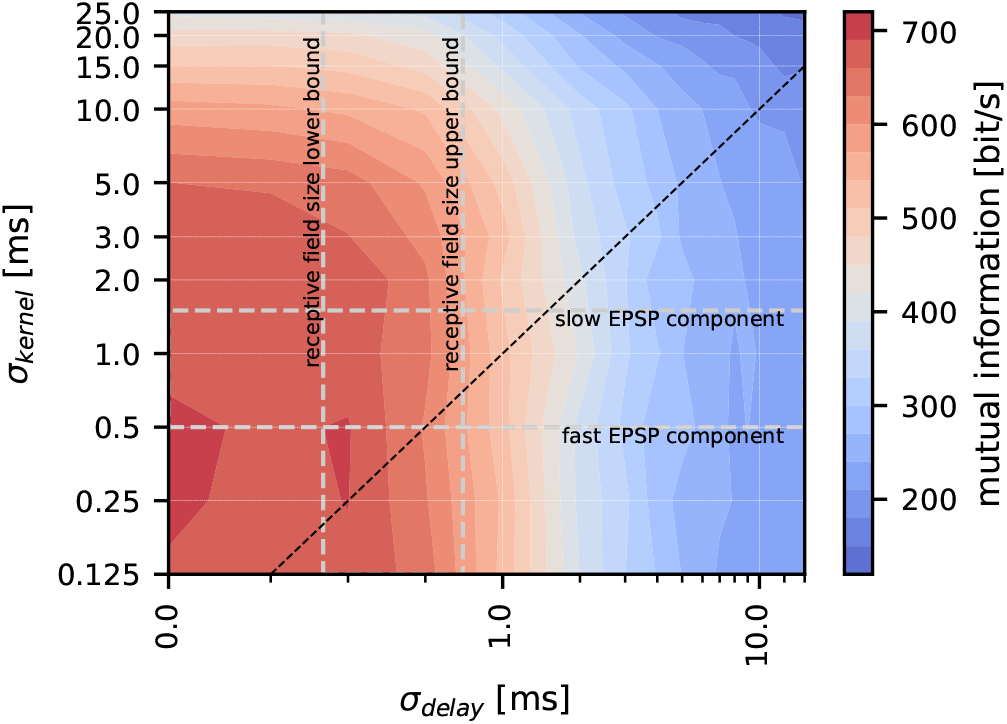
Information filtering by conduction delays is stronger than by synaptic filtering. Comparison of the effects of conduction delays (x-axis) and synaptic transfer (y-axis) on the information content of the population response (color code). Dashed black line is the identity line. Area between horizontal dashed lines depicts the standard deviations (*σ_kernel_*) of the fast and slow EPSP components as reported for the P-unit to pyramidal cell synapses [55]. Vertical dashed lines enclose the range of conduction delays that can be expected for pyramidal neurons in the lateral segment of the electrosensory lateral line lobe [29] considering the 2σ or 4σ receptive field sizes. The data shown here assumes a population size of 16 cells.

In *A. leptorhynchus* P-units project onto pyramidal cells in three adjacent maps of the ELL [17, 25, 29] which have different receptive field sizes and spectral tuning properties [28, 29]. Receptive fields are largest in the lateral segment where pyramidal cells integrate about 1000 P-units [29]. Furthermore, from studies on the EPSPS in the ELL we can estimate the time-constants of synaptic filtering [55]. The horizontal and vertical dashed lines in Fig 7 depict lower and upper estimates of the receptive field sizes and synaptic kernels in the lateral segment. The intersection area highlights the real combination. It appears that especially the receptive field size is a good compromise for maximizing the gain from population coding and minimizing the loss due to conduction delays. The current view on the ELL is that lateral segment cells encode high-frequency global signals as arise from social interactions while, on the other extreme, the cells in the centro-medial segment with small receptive fields (about 10 P-units), and higher sensitivity for low-frequency signals are devoted to the processing of navigation and prey-related signals. Confusingly, small receptive fields would be particularly well suited to encode high frequency signals as attenuation by conduction delay spread is minimal. It was previously found that P-units not only encode the AM in their firing rate but that spikes simultaneously carry precise timing information [40]. Timing information was found to be conserved in pyramidal neurons in the lateral segment, though to a lesser extent than in the input layer. Future studies should address whether smaller receptive fields in the centro-medial conserve timing information better than the larger receptive fields in the lateral segment.

### Local computations or fast conduction to preserve timing

The fish’s electrosensory surface is very large and our results suggest that convergence is limited to maintain a balance between information gain through convergence and spread of conduction delays and thus the dynamic range of signal encoding. What about other sensory surfaces? The olfactory systems of vertebrates and invertebrates share similarities olfactory receptor neurons (ORNs) are tuned to certain odorants and project to the olfactory bulb, respectively the antennal lobe. There, all similarly tuned neurons converge onto interneurons (projection neurons, PNs) that are organized in Glomeruli [56]. Commonly, olfactory stimulus dynamics are considered to be slow. In odor plumes, however, stimulus intensities may vary considerably on a millisecond timescale [57] and olfactory receptor neurons actually encode such dynamic changes of odor concentrations [58]. Preserving such time-critical information may require rather local computations.

## Conclusion

In their great book on the principles of neural design Sterlin and Laughlin work out three main principles that define the way nervous systems should be designed [59]: (i) Only the absolutely required information should be sent down axons, (ii) sending information should employ as few action potentials as possible and (iii) the system should be designed to minimize the use of axons in terms of length and diameter. Here we would like to add considerations about the compromise of encodable stimulus dynamics, receptive field size, and information (i.e. the signal to noise ratio).

Reading out information from a sensory surface must be a compromise of the receptive field size (the population size) and the signal dynamics that need to be encoded. This again emphasizes the neuroethological claim that we can only understand the brain when we consider it within its natural context with knowledge about the natural stimuli and the extracted features [60–63].

## Supporting information

**S1 Fig.**
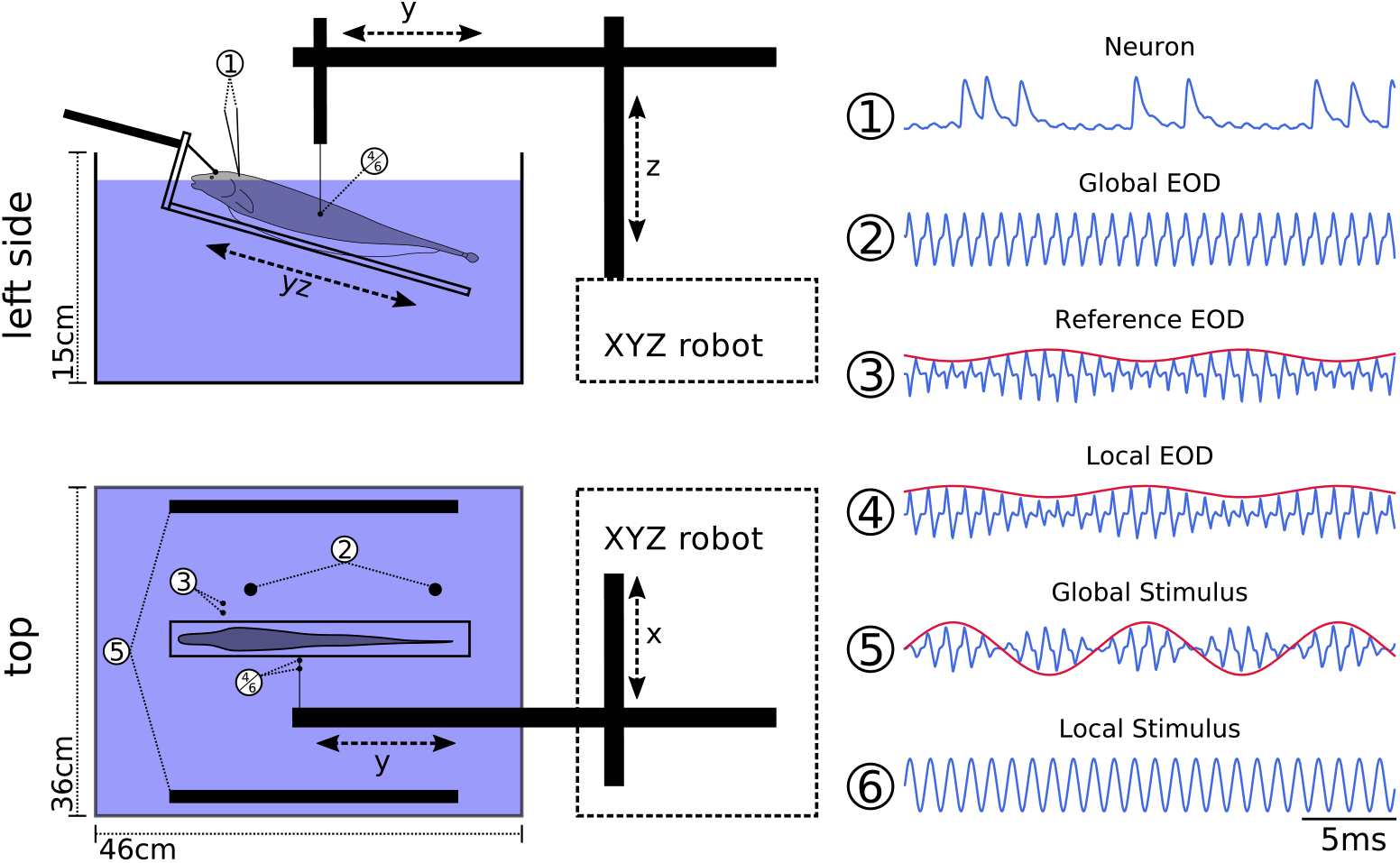
Experimental setup. Schematics of the side and top view of the experimental tank. Numbers refer to the recorded signals depicted on the right. **1:** The semi-intracellular potential was recorded in the lateral line nerve. **2–4** The electric field of the fish was recorded in three different ways. From top to bottom; **2:** *Global EOD* head to tail measurement, measurement electrodes were placed isopotential to the stimulus electrodes to record the unperturbed field of the fish. **3:** The *Reference EOD* is a proxy of the transdermal potential picked up by the electroreceptor afferents and is measured using a pair of silver wired oriented orthogonal to the body axis of the fish and placed just posterior of the operculum. **4:** Local EOD measurement dipole mounted on the robot arm. Via two carbon rods (**5**) the global stimulus could be given. A local stimulus was given via the dipole electrode **6**.

**S2 Fig.**
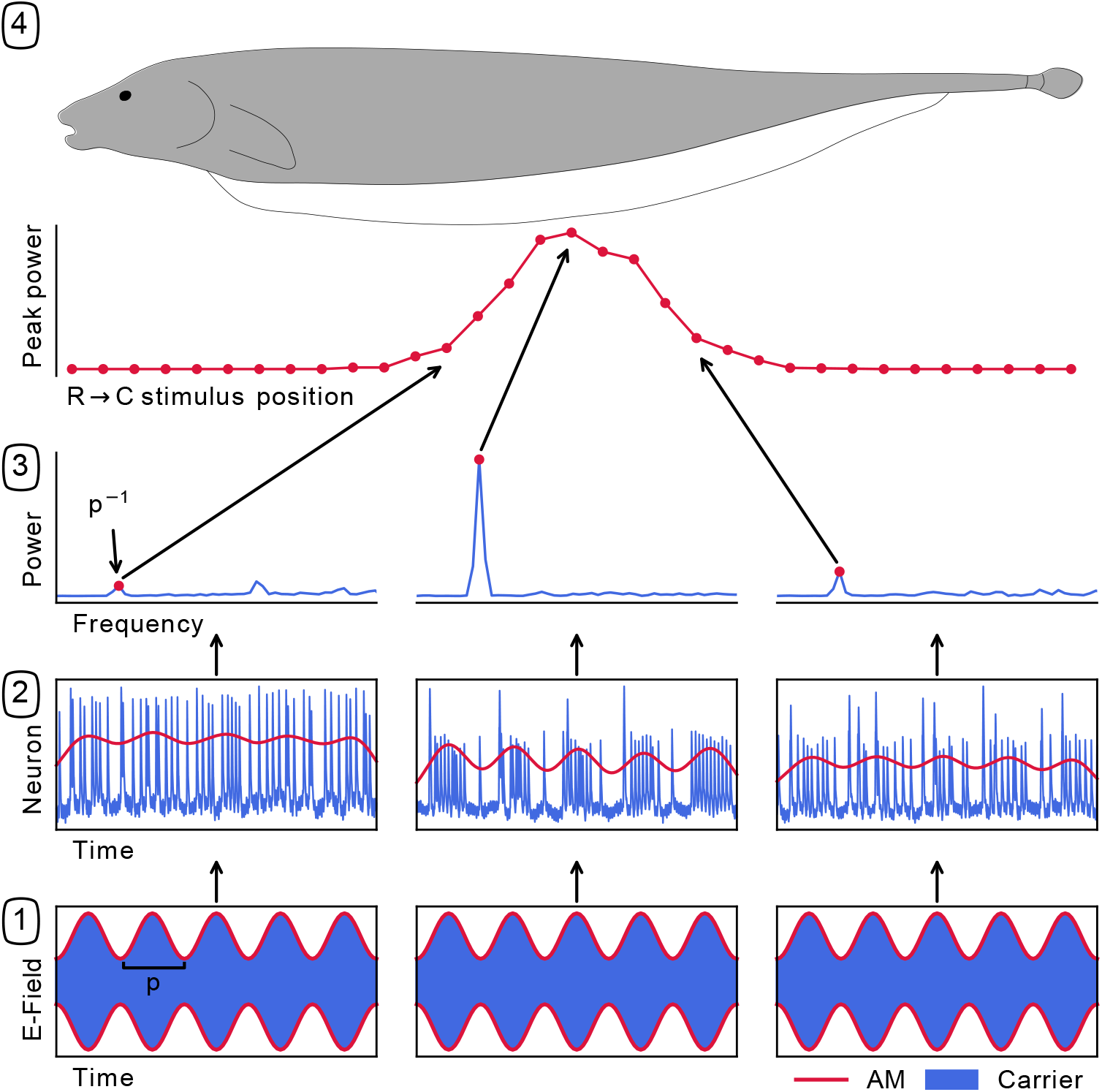
Estimating receptor position. The location of a recorded P-unit on the body of the fish was estimated by moving the local stimulus dipole alongside the rostro-caudal axis of the animal. At each position a stimulus was presented that led to an amplitude modulation of the recorded fish’s EOD (bottom trace, the blue line is the carrier, i.e. the fish’s EID, the red line indicates the induced amplitude modulation with the period p). The neuronal spiking response (panels in row 2) were measured and the firing rate was estimated unsing kernel convolution with a Gaussian kernel (red line). The power at the expected frequency *p*^−1^ was extracted from the power spectra of the firing rate (panels in row 3) and was then plotted as a function of the rostro-caudal position (panel 4). The receptor location was the maximum position of a fitted Gaussian.

**S3 Fig.**
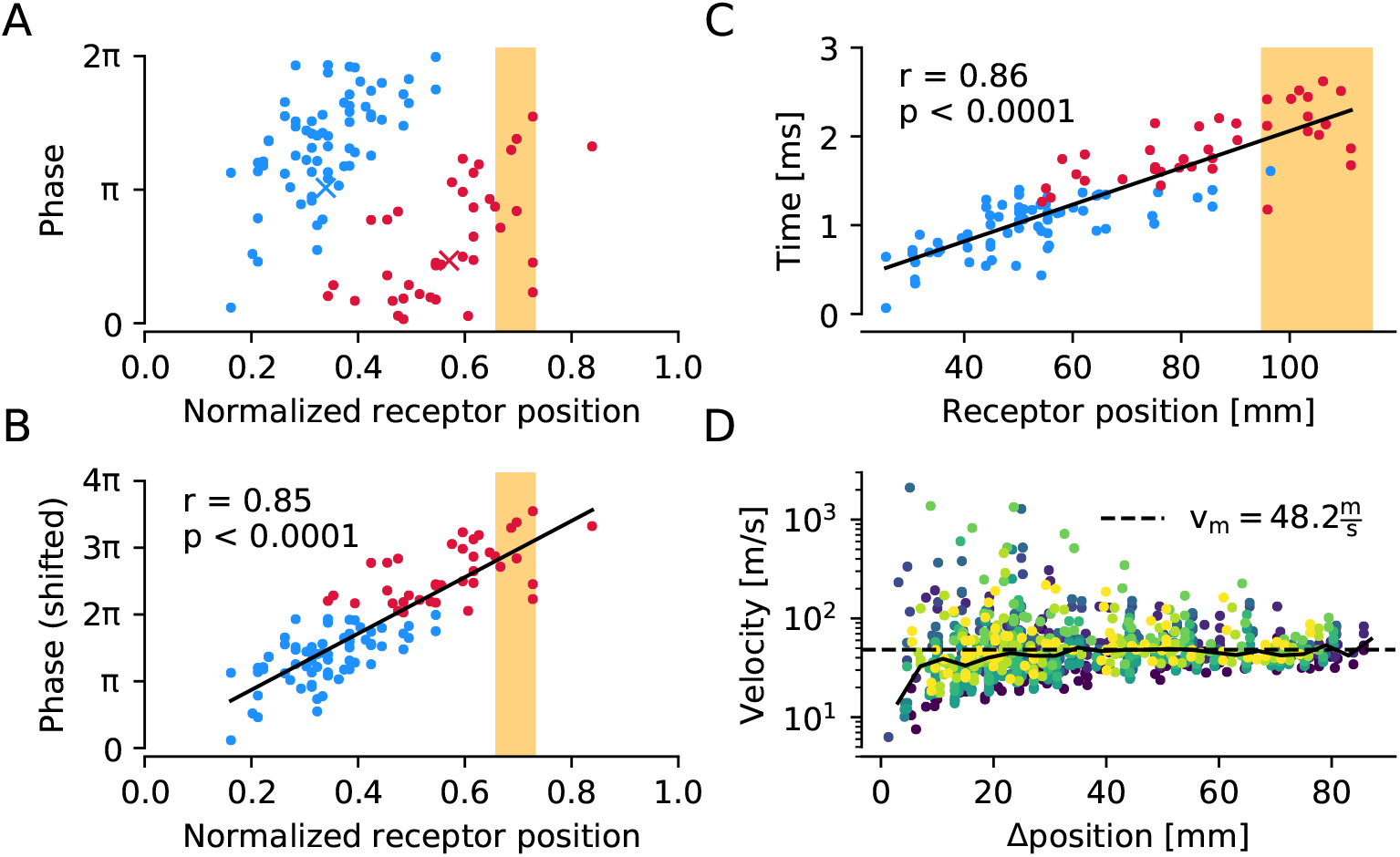
Dependency of phase locking on receptor position. **A:** Preferred phase relation between the baseline spikes and the fish’s own EOD measured at the operculum (*Reference EOD*, S1 Fig. Experimental setup). Colors represent clusters as established by K-means clustering. Crosses depict cluster centroids. Orange section shows the region in which the sign of the EOD is inverted (*Local EOD*). **B:** The red cluster was phase shifted by one cycle (2π). Now phases correlate positively with the rostro-caudal receptor position (solid line, Pearson correlation). **C:** Replotting of the data in absolute units. Spike phases are expressed as the absolute delay between the beginning of the EOD (ascending zero crossing) and the respective action potential. The receptor positions on the rostro-caudal axis are given in absolute numbers relative to the snout position. The slope of the regression line is the inverse of the conduction velocity of 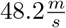. **D:** Conduction velocity estimated from all possible pairs of P-units recorded in the same animal. For each pair the conduction velocity is estimated from the difference in phase relation to the EOD and the difference in rostro-caudal position. Colors indicate cell identity of the reference cell against which velocity differences between cells were measured. Dashed line is the conduction velocity expected from **C** and the solid line connects the medians estimated for the pairs. There is a good agreement between the two approaches chosen in B and D with less variability for larger distances between cells in the pairs.

**S4 Fig.**
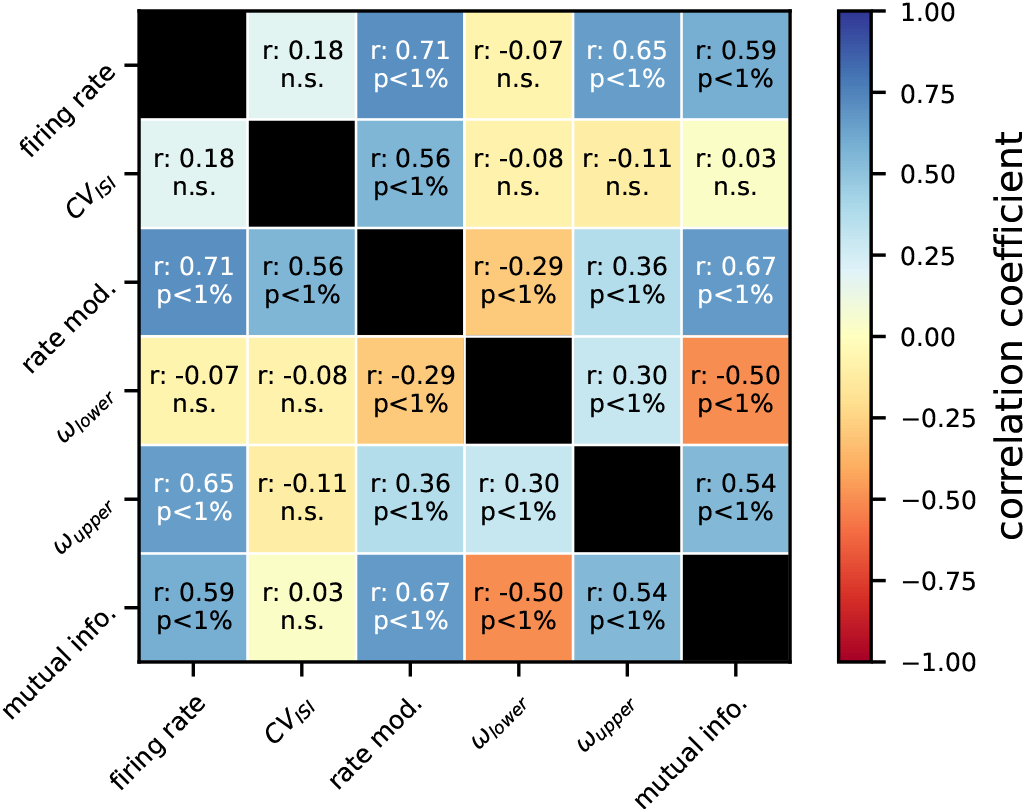
Correlations of cellular response properties. Color codes for the Pearson correlation coefficient ‘r’ with blue colors depicting positive and red colors depicting negative correlations. p-values are Bonferroni corrected (n=15 correlations). A statistically significant correlation was assumed for p < 0.05.

**S5 Fig.**
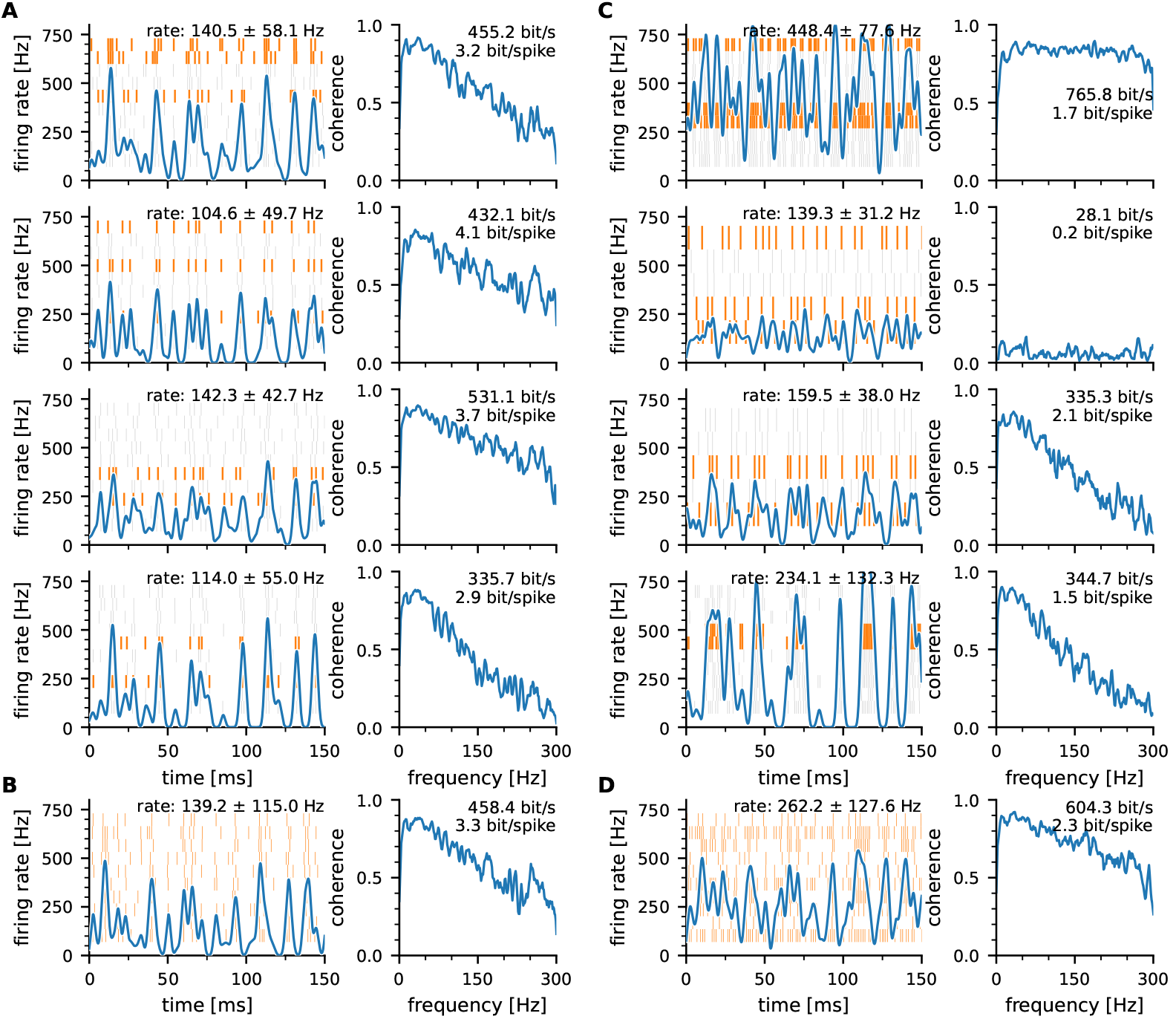
Stimulus encoding with homogeneous and heterogeneous populations. **A:** Responses of four example cells to the same white noise stimulus. The cells were chosen based on similar the average firing rates (100 to 15 Hz) and similar response modulations (40 to 60 Hz). Left: The raster plot in the background depict the spike times of up to 10 consecutively recorded stimulus repetitions (150 ms out of 10 s total trial duration). Orange trials were randomly selected to create the population response shown in B. Blue line depicts the across-trial firing rate estimated by kernel convolution with a Gaussian kernel with a standard deviation of 1.25 ms. Right: stimulus response coherence smoothed with a five point running average and based on segments of 16384 samples (0.82 s) and 50% overlap. Mutual information is calculated according to eq. 8. **B:** Population response of a “homogeneous” population created as the average of the orange-highlighted trials in A. **C:** Same as A, but for four example cells with different response properties. Maximum and minimum firing rate (top and second row) minimum and maximum response modulation (third and fourth row). **D:** Same as B but trials were selected from the heterogeneous cells shown in C.

## Acknowledgments

We thank Jan Benda, Felix Franke, Benjamin Lindner, and Greg Knoll for critical reading and in-depth discussions. We further acknowledge support by the Open Access Publishing Fund, University of Tübingen.

